# Connecting the Dots: Approaching a Standardized Nomenclature for Molecular Connectivity Combining Data and Literature

**DOI:** 10.1101/2024.05.10.593490

**Authors:** MB Reed, L Cocchi, GM Knudsen, C Sander, G Gryglewski, J Chen, T Volpi, P Fisher, N Khattar, LR Silberbauer, M Murgaš, GM Godbersen, L Nics, M Walter, M Hacker, A Hammers, TR Ogden, JJ Mann, B Biswal, B Rosen, R Carson, J Price, R Lanzenberger, A Hahn

## Abstract

PET-based connectivity computation is a molecular approach that complements fMRI-derived functional connectivity. However, the diversity of methodologies and terms employed in PET connectivity analysis has resulted in ambiguities and confounded interpretations, highlighting the need for a standardized nomenclature.

Drawing parallels from other imaging modalities, we propose “molecular connectivity” as an umbrella term to characterize statistical dependencies between PET signals across brain regions at the individual level (within-subject). Like fMRI resting-state functional connectivity, “molecular connectivity” leverages temporal associations in the PET signal to derive brain network associations. Another within-subject approach evaluates regional similarities of tracer kinetics, which are unique in PET imaging, thus referred to as “kinetic connectivity”. On the other hand, “molecular covariance” denotes group-level computations of covariance matrices across-subject. Further specification of the terminology can be achieved by including the employed radioligand, such as “metabolic connectivity/covariance” for [^18^F]FDG as well as “tau/amyloid covariance” for [^18^F]flutemetamol / [^18^F]flortaucipir.

To augment these distinctions, high-temporal resolution functional [^18^F]FDG PET scans from 17 healthy participants were analysed with common techniques of molecular connectivity and covariance, allowing for a data-driven support of the above terminology. Our findings demonstrate that temporal band-pass filtering yields structured network organization, whereas other techniques like 3^rd^ order polynomial fitting, spatio-temporal filtering and baseline normalization require further methodological refinement for high-temporal resolution data. Conversely, molecular covariance from across-subject data provided a simple means to estimate brain region interactions through regularized or sparse inverse covariance estimation.

A standardized nomenclature in PET-based connectivity research can reduce ambiguity, enhance reproducibility, and facilitate interpretability across radiotracers and imaging modalities. Via a data-driven approach, this work provides a transparent framework for categorizing and comparing PET-derived connectivity and covariance metrics, laying the foundation for future investigations in the field.

## Introduction

The assessment of resting-state functional brain networks, as mostly elucidated through functional Magnetic Resonance Imaging (fMRI) and Electroencephalography (EEG), has been a cornerstone of neuroimaging research for decades due to its low risk, low cost, and widely available hard- and software. Resting-state functional connectivity (FC) has provided valuable insights into the organization of the brain and network interactions by correlating moment-to-moment fluctuations of signals between spatially distinct brain regions at rest. Positron Emission Tomography (PET) enables imaging of physiological processes at the molecular level, capable of detecting energy metabolism, neuronal receptors, enzymes, and other targets at nanomolar concentrations. However, its application in connectivity analyses remains relatively unexplored. While molecular connectivity is a concept dating back to the 1980s ^1^ and 1990s ^2,3^, little progress has been made, in part due to technological constraints in PET imaging that resulted in limited count rates at high temporal resolutions. These constraints precluded the reconstruction of dynamic PET data in the range of seconds and thus, the estimation of connectivity at the individual level. Consequently, and due in part to its simplicity, the computation of covariance (i.e., not in a statistical sense) metrics across subjects remained the commonest approach as a proxy for *molecular* connectivity. The widespread availability of [^18^F]Fluorodeoxyglucose ([^18^F]FDG) for *metabolic* connectivity (i.e., molecular connectivity for glucose metabolism), represents a promising avenue for probing brain interactions based on metabolic demands, complementing its fMRI counterparts ^4–8^. However, the inherent disadvantage of estimating associations across an entire group of subjects instead of connectivity at the subject level is a major obstacle regarding its individual biological interpretation ^9,10^. Figure 1 presents a graphical overview of common techniques used to assess brain connectivity in humans *in vivo*.

**Figure 1:**
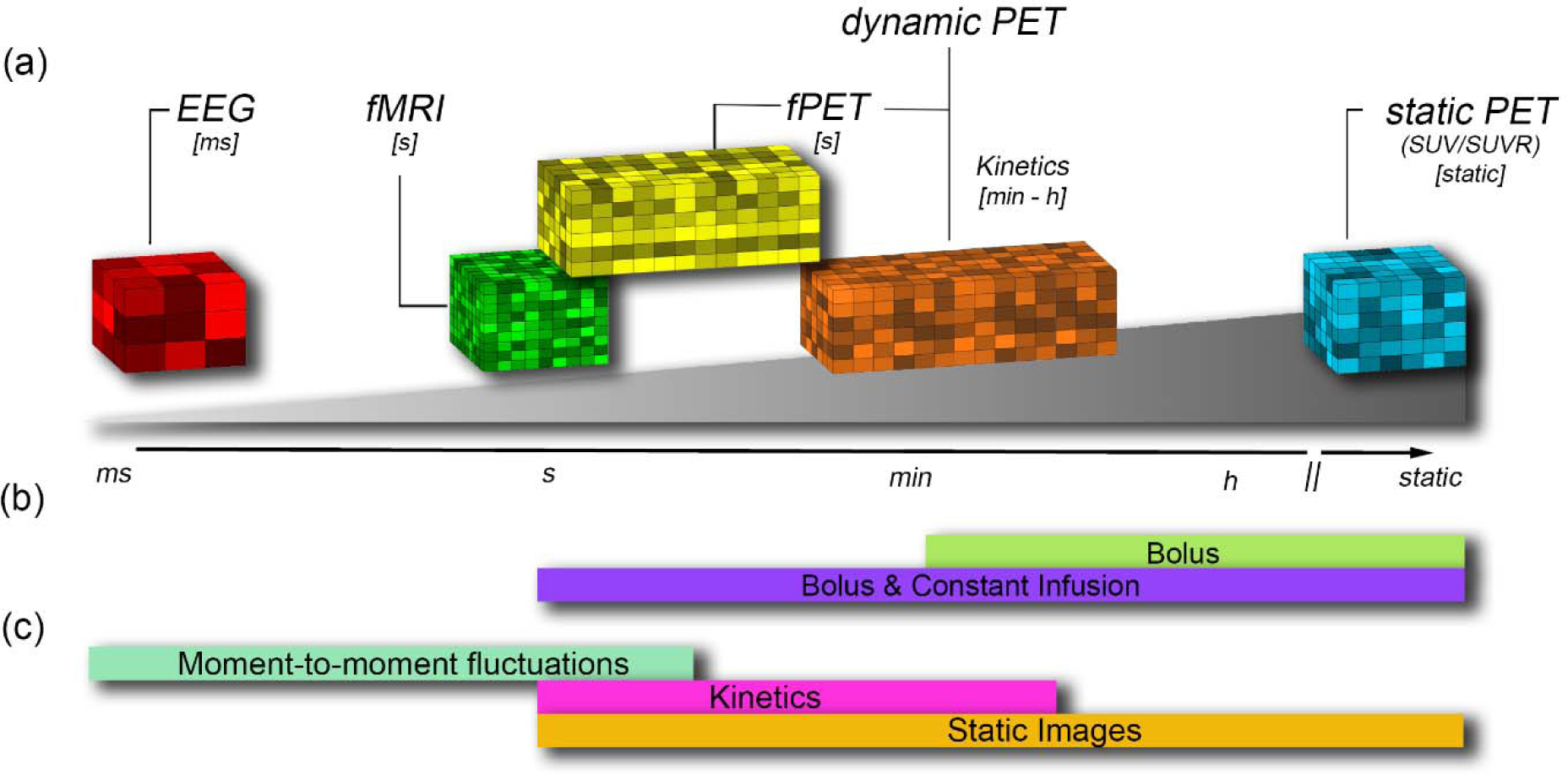
Graphical overview of common neurophysiological techniques used to assess brain connectivity. (a) Acquisition: Each voxel represents a common imaging method where the position on the x-axis indicates the temporal resolution of each technique as an input for different connectivity measures. The voxel itself is representative of the spatial resolution of the technique. (b) Radiotracer administration: In the field of PET imaging, application of the radiotracer as bolus enables to obtain static images and tracer kinetics, while a bolus+constant infusion protocol additionally allows to capture moment-to-moment signal fluctuations. (c) Input for connectivtiy estimation: Dynamic approaches enable the computation of within-subject connectivity, which is commonly calculated by correlating the time courses of amoung brain regions. For PET this is derived from either bolus plus constant infusion or simple bolus. In contrast, across-subject covariance is estimated over a group of participants as it lacks the temporal component. As such, these techniques use different input signals to estimate connectivity, (i.e., moment-to-moment fluctuations in the signal or radiotracer kinetics) and across-subjects PET signal covariance (i.e., static images). Furthermore, static images can also be obtained from dynamic PET data, e.g., through kinetic modeling, and subsequently enable estimaton of covariance metrics. EEG: electroencephalography, fMRI: functional magnetic resonance imaging, fPET: functional positron emission tomography, SUV(R): standardized uptake value (ratio).

Recent technological progress has transformed the landscape of molecular connectivity research. With increased sensitivity PET scanners, standardized infusion protocols (i.e., bolus + constant infusion), refined reconstruction algorithms, and advanced pre-processing including filtering techniques and post-processing, researchers are now better equipped to investigate brain networks on a molecular level. These advances allow using PET data at previously unprecedented temporal resolutions within the range of minutes and seconds ^9,11,12^, more akin to that of fMRI ^11^. This has established the foundation for estimating individual temporal molecular connectivity through various computational methodologies. These include the application of within-subject Euclidean distance metrics ^13^, a third-order polynomial function ^14,15^, spatiotemporal filters ^9^, as well as the utilization of across-subject covariance matrices ^16–19^ as well as hybrid approaches to integrate fMRI and PET metrics ^20,21^. Most of these approaches aim to compute connectivity at an individual level by using temporal information from the PET data. Furthermore, the 3^rd^ order polynomial, spatiotemporal filter and baseline normalization aim to correlate moment-to-moment fluctuations in the PET signal, while the Euclidian distance evaluates differences in tracer kinetics. Exceptions to this are covariance matrices and sparse inverse covariance estimation (SICE), which compute associations between brain regions across a group of subjects.

Unfortunately, each technique has been labelled as molecular connectivity, despite differences in the underlying assumptions, computations, and outcome metrics, resulting in ambiguous terminology. Moreover, related terms such as “metabolic connectivity mapping” are employed to describe various outcomes, leading to potential confusion ^13,20^.

As the field experiences a growth in utilization and methodological diversity, there is a pressing need for standardization in nomenclature. The absence of a unified terminology poses challenges in synthesizing findings across studies and impedes the establishment of a cohesive framework for interpreting PET connectivity outcomes. Discussions regarding the definition of molecular connectivity and covariance, as well as the distinct yet valuable insights offered by each approach, have already commenced ^9,10,18^. However, previous work either compared only a subset of approaches or was qualitative in nature, while widespread consensus grounded in the actual outcome parameters of each technique is missing.

We aim to address this gap by proposing a standardized nomenclature for the most utilized PET connectivity techniques across a multidisciplinary, international group of researchers in the field. Our proposed nomenclature is based on a comprehensive review of existing literature on molecular connectivity and covariance techniques (part 1: nomenclature). To validate this proposal, we conducted a showcase using high-temporal resolution [^18^F]FDG data, which was previously unavailable for such analyses (part 2: experimental data). This approach ensures that our terminology not only aligns with established methodologies but also demonstrates practical feasibility in an experimental setting. By integrating theoretical foundations with empirical comparisons, we establish a cohesive and robust framework for defining molecular connectivity metrics.

### Nomenclature of molecular connectivity and covariance

The standardization of nomenclature for quantification of PET radioligands ^22^ has significantly enhanced the clarity of subsequent research, fostering greater consistency in findings. Given the substantial growth in interest in PET connectivity research in recent years, the standardization of its nomenclature is fundamental, as it addresses several pivotal issues within the field. Firstly, the utilization of a single term to describe multiple distinct concepts (and vice versa) can cause confusion among researchers, impeding the interpretation of study outcomes. By harmonizing terminology, researchers can facilitate seamless communication and collaboration, thereby increasing the reproducibility of findings across studies. Additionally, a standardized nomenclature enhances comprehension, particularly among diverse audiences and within broader contexts. Neuroimaging traverses’ various disciplines and encompasses researchers with diverse levels of expertise. Standardized terminology enables researchers from disparate backgrounds to readily compare findings, fostering knowledge exchange and interdisciplinary cooperation.

Historically, the grouping of PET connectivity methodologies has delineated the term ℌmolecular connectivityℍ (and with the common use of [^18^F]FDG also “metabolic connectivity”) as an umbrella concept encompassing both within-subject connectivity and across-subject metrics. However, the interchangeable use of ℌmolecular connectivityℍ for within-subject and across-subject associations is problematic, as these represent fundamentally distinct measures. Most imaging modalities use “connectivity” to characterize the strength (and potential directionality) of couplings between brain regions within individuals (Figure 1). This is true for functional connectivity obtained from fMRI and EEG data and structural connectivity derived from diffusion-weighted MRI. On the other hand, “covariance” metrics gauge the statistical associations between regions of interest of a static outcome metric across individuals, such as gray matter volume obtained from T1-weighted structural images ^23,24^ or SUVR ^25^ in PET imaging. A clear differentiation between these concepts is imperative to prevent misinterpretation and ensure accurate communication of study findings (Figure 2).

**Figure 2:**
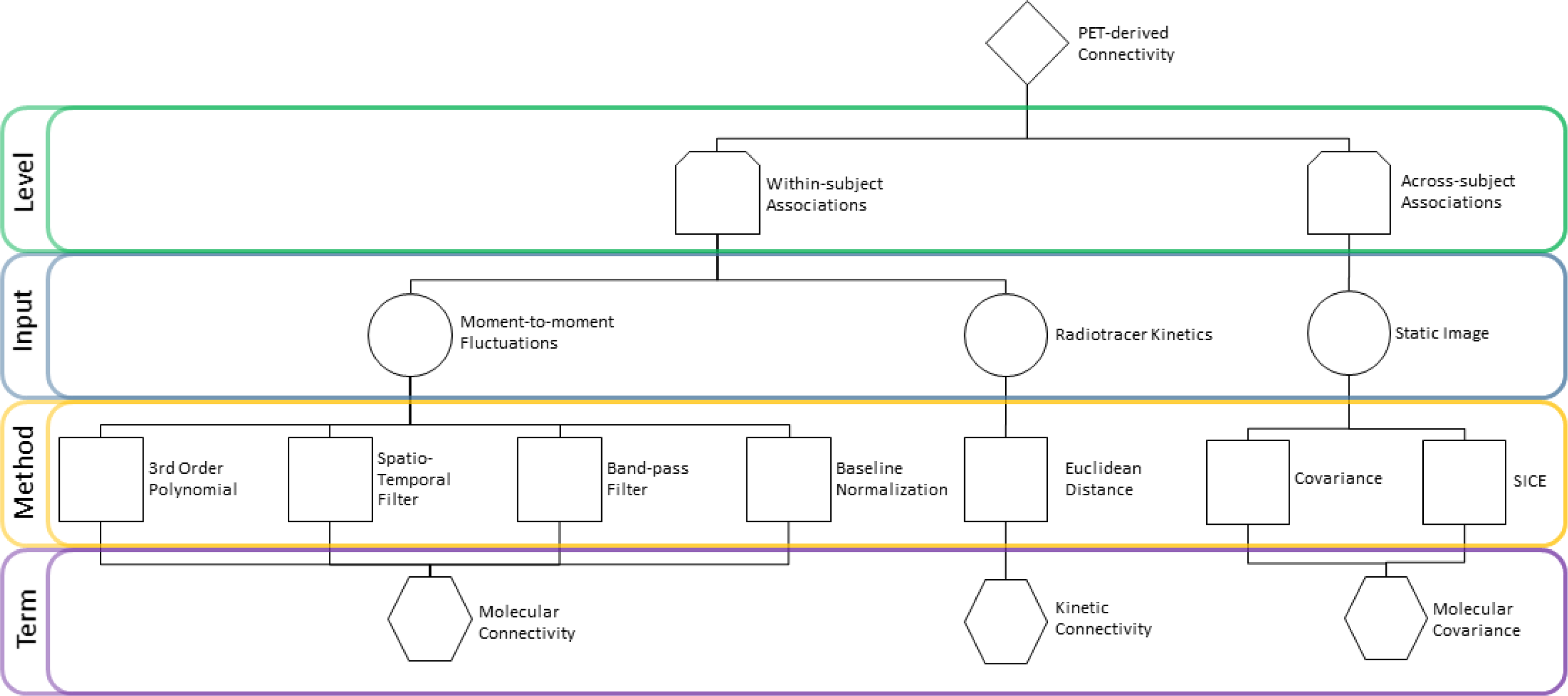
Summary of techniques and proposed nomenclature of PET-based connectivity. Within-subject connectivity is broadly separated into the assessment of moment-to-moment signal fluctuations and radiotracer kinetics. The former requires the elimination of the low-frequency baseline radiotracer uptake and reducing high-frequency noise, which a band-pass filter can achieve. In their current form, the 3^rd^ order polynomial and spatio-temporal filter were not able to remove noise, and the baseline normalization still included pronounced baseline radiotracer uptake. Despite their different preprocessing, correlation of moment-to-moment signal fluctuations was used for all approaches to estimate “molecular connectivity”, which is also most alike to fMRI-based functional connectivity. On the other hand, the Euclidian distance of time activity curves identifies differences in radiotracer kinetics between brain regions, thus termed “kinetic connectivity”. The across-subject metrics use static images as input and after the computation of covariance and SICE yield “molecular covariance”

For “molecular connectivity” analyses, we therefore suggest that this term may encompass all approaches that use moment-to-moment fluctuations in the PET signal to estimate connectivity (like “functional connectivity” used in fMRI). In this context, further detail emerges for *within-subject* connectivity methods, each capturing *temporal dynamics within an individual*, contingent upon the method employed to estimate connectivity metrics. On the other hand, “molecular covariance” should be used for the estimation of network interactions *across-subjects*. The discourse on the disparities between molecular connectivity and molecular covariance is not novel ^9,10,26^, and will therefore not be repeated here. Nevertheless, these discussions underscore the relevance of *within-subject* and *across-subject* connectivity metrics in PET connectivity research and their inherent differences. While within-subject connectivity furnishes insights into individual variability and dynamic network properties, across-subject covariance offers a broader perspective on shared connectivity patterns across populations. Another PET-specific approach distinct from all former methods is the Euclidean distance calculation. This divergence stems from the diversity of inputs it accommodates (raw or compartment specific TACs) and how connectivity is calculated (Euclidean distance between TACs). In this regard, the method is theoretically limited to estimating solely positive connectivity metrics. The assumptions and calculations of this technique are based on the kinetics of the radiotracer across an entire PET scan, leading us to propose the term “kinetic connectivity” for a clear distinction of this PET approach. Notably, extension with kinetic modelling allows to derive kinetic connectivity for the individual compartments, yielding relevant information regarding the separation of the transport across the blood-brain barrier (K_1_, k_2_) and irreversible uptake into the cells (k_3_). The proposed terminology also reflects these distinct effects.

Regarding different radiotracers, we would like to propose that the term “molecular connectivity/covariance” serves well as an umbrella term for PET-based estimation of network interactions in general. This also leaves ample opportunity for further specification of different radioligands and targets such as “metabolic connectivity/covariance” when using [^18^F]FDG, “5-HT_1A_ covariance” for [*carbonyl*-^11^C]WAY100635 ^27^, “SERT covariance” for [^11^C]DASB ^28^ or “tau/amyloid covariance” for [^18^F]flutemetamol (amyloid-β) / [^18^F]flortaucipir (tau) ^29^ (see supplementary table 1 for a list of examples). Likewise, the term “kinetic connectivity” can be extended for the employed radiotracer and/or target, such as “metabolic kinetic connectivity,” when using [^18^F]FDG. We expect that this field will experience rapid growth and application to other target structures in the near future, further underlining the need for standardized terminology and assessment of the feasibility and interpretation for radioligands beyond [^18^F]FDG.

Recently, also hybrid connectivity and covariance techniques have emerged, leveraging the complementary strengths of static PET and fMRI-based functional connectivity to delineate molecular covariance ^21^ or estimate directional connectivity ^20^. Notably, recent advancements have also integrated dynamic fPET with fMRI to investigate task-related neuronal responses ^30^. Further extension of such combined analyses to connectivity and exploration of hybrid methods, which integrate data from multiple modalities such as fPET, fMRI and EEG, holds promise for providing a comprehensive understanding of brain connectivity. These hybrid approaches could also enable the investigation of directional connectivity (such as dynamic causal modeling for fMRI data), shedding light on the causal interactions between brain regions and enhancing our ability to elucidate complex neural networks underlying cognition and behavior. As the field expands, the agreement and adoption of specific terms will become increasingly necessary, see Table 1 for an overview of the proposed terms and their definitions.

**Table 1:** Description of each PET connectivity and covariance term and its respective abbreviation, level of estimation and definition.

| Term | Abbreviation | Level | Definition |
| --- | --- | --- | --- |
| PET-based Connectivity |  |  | PET-based connectivity refers to the assessment of connections and interactions between molecular entities within the brain. It is an umbrella term that encompasses all subsequent terms. |
| Molecular Connectivity | MC | Within-subject | Molecular connectivity estimates associations between brain regions at the individual subject level, e.g. by correlation of moment-to-moment signal fluctuations. This is akin to functional connectivity obtained with fMRI. |
| Kinetic Connectivity | KC | Within-subject | Kinetic connectivity evaluates the similarity of radiotracer kinetics across various brain regions, as identified through time-activity curves within an individual. |
| Molecular Covariance | MCov | Across-subject | Molecular covariance utilize PET-derived static images (SUV/SUVR, binding potential, etc.) across subjects to estimate interregional associations within the brain. This is akin to structural covariance obtained with T1-weighted MRI. |

### Experimental data

In this chapter we aim to augment the literature-based nomenclature by comparing established methodologies for estimating molecular connectivity and molecular covariance using high-temporal resolution [^18^F]FDG fPET data at resting-state alongside their fMRI-derived functional connectivity counterparts. We aim to show the similarities and differences between these methodologies, thereby providing a data-driven standardization of PET connectivity nomenclature.

### Participants and demographics

Seventeen healthy volunteers (mean age ± SD = 24.6 ± 3.5 ranging from 21 to 32 years old, 10 female) were recruited for the fPET molecular connectivity analysis, while 62 healthy participants (mean age ± SD = 29.0 ± 9.25 ranging from 18 to 51 years old, 35 female) were recruited for the fMRI functional connectivity analysis.

Participants’ general health status was evaluated through a comprehensive medical evaluation, including medical history, physical examination, electrocardiogram, and routine laboratory tests. The Structured Clinical Interview for DSM-IV for Axis I disorders (SCID-I) was employed to exclude any prior or current psychiatric disorders. Exclusion criteria encompassed chronic medical conditions, psychiatric disorders, current or past substance use disorder, psychopharmacological treatment, and contraindications for PET/MR scans, such as implants, claustrophobia, and research-specific radiation exposure. Urine drug tests were conducted during screening, and pregnancy tests were administered to female participants both at screening and before each scan. All participants provided written informed consent and received financial compensation for their involvement.

This study was approved by the ethics committee of the Medical University of Vienna (EK 1307/2014) and adhered to the principles outlined in the Declaration of Helsinki. This investigation constitutes a component of a larger preregistered study (clinicaltrials.gov, NCT02711215).

### Study design

For the fPET acquisition, each participant underwent two 50-minute hybrid [^18^F]FDG fPET/fMRI scans. Following a baseline acquisition of 20 minutes at resting-state, a double-blind pharmacological challenge of either citalopram 8 mg or saline was administered. While performing connectivity analyses in the current project, data between 5 (i.e. after radiotracer uptake is in a steady state for bolus + infusion ^31,32^) and 20 minutes were extracted from the first measurement, i.e. before any drug application occurred. This allowed the participants to be in the same cognitive state (here: resting-state) during the experiment.

fMRI acquisition was carried out in a similar manner and using the same scanner system, though in different individuals as described previously ^33^. Here, 10-minute resting-state fMRI data from the placebo measurement were utilized, a more detailed description of the study design can be found in ^33^.

### Data acquisition

Participants were instructed to fast, except for water intake, for a minimum of 5.5 hours preceding the administration of the radioligand ^34^. A cannula was inserted into the radial artery for arterial blood sampling (which was not used in this work), while two cannulas were placed in a cubital vein of the contralateral arm for the infusion of the radiotracer [^18^F]FDG and study medication (citalopram or placebo). Synthesis of [^18^F]FDG was conducted following a well-established protocol ^35^. The radiotracer (5.1 MBq/kg) was administered as a bolus (1020 kBq/kg/min, 1 minute) followed by a continuous infusion (83.3 kBq/kg/min, 49 minutes) utilizing a perfusion pump (Syramed µSP6000, Arcomed, Regensdorf, Switzerland) situated within an MR-shielded environment (UniQUE, Arcomed). All scans were conducted using a hybrid 3T PET/MRI scanner (mMR Biograph, Siemens Healthineers, Germany).

Resting-state fMRI data was acquired for 10 min using an echo-planar imaging sequence (TE/TR= 30/2440 ms, 2.1 × 2.1 mm in-plane resolution, 3 mm slice thickness with 0.75 mm gap), 100 × 100 voxels in-plane, 36 slices, GRAPPA 2.

Before radiotracer administration, a structural image was obtained using a T1-weighted MPRAGE sequence with the following parameters: TE/TR = 4.21/2200 ms, TI = 900 ms, flip angle = 9°, matrix size = 240×256, 160 slices, voxel size = 1 × 1 × 1 mm + 0.1 mm gap, TA = 7:41 min, which was used for spatial normalization of the fPET data. fPET acquisition commenced subsequently, following the protocol described above and in previous work ^12^.

### fMRI preprocessing and filtering

fMRI data were preprocessed in SPM12 and ArtRepair toolbox ^36^, following an established protocol ^37^. In brief, the data underwent correction for transient slice artifacts and slice-timing discrepancies, followed by realignment and reslicing of the realigned images. Pre-smoothing with a 4 mm FWHM Gaussian kernel was applied, followed by motion regression, detection of motion outliers, and subsequent regression ^38^ followed by despiking ^36^. Subsequently, the data were normalized to MNI space with an isotropic resolution of 2 mm ^3^. Acknowledging the substantial variability in fMRI connectivity patterns across preprocessing steps, including common corrections for cerebrospinal fluid and white-matter signals in resting-state fMRI data, we intentionally omitted such corrections. This decision was made to ensure consistency in data preprocessing across modalities. Data were band-pass filtered (0.01-0.1 Hz) ^39,40^.

### fPET preprocessing

fPET image reconstruction and preprocessing followed established protocols outlined in our previous publications ^11,12,41^, keeping the processing as similar as possible to fMRI. List-mode PET data underwent reconstruction into frames of 3 seconds utilizing an ordinary Poisson-ordered subset expectation maximization algorithm (3 iterations and 21 subsets, OP-OSEM), with a matrix size of 344 × 344 and 127 slices with a voxel size of 2.09 × 2.09 × 2.03 mm and a NEMA resolution of 4.3 mm ^42^. Attenuation and scatter correction were performed using a pseudo-CT approach based on the structural T1-weighted image ^43^. fPET data were preprocessed utilizing SPM12 (Wellcome Trust Centre for Neuroimaging) and included head movement correction (quality = best, registration to mean image) and coregistration to the structural image. The structural MRI was spatially normalized to the standard space defined by the Montreal Neurological Institute (MNI), and the transformation matrix was applied to the coregistered fPET data. Spatial and temporal smoothing was performed using a dynamic non-local means (NLM) filter with a search window of D = 11 voxels and a patch size of 3 × 3 × 3 voxels and 5 frames ^44,45^, followed by Gaussian smoothing with a FWHM of 5 mm ^11^. Compared to a standard 3D Gaussian filter, this approach has the advantage of increasing the signal-to-noise ratio while enabling the capture of acute temporal changes in the signal ^46^.

### Connectivity and covariance estimation

To distinguish between the different type of connectivity and covariance methods, within-subject measures are prefixed with (A) and across-subject with (B).

#### (A1) 3^rd^ order Polynomial

The application of a 3^rd^ order polynomial fitting is employed to model the cumulative behavior of [^18^F]FDG uptake extracted from the time course of each brain region. This method, as outlined by ^14,15^, aims to capture the temporal dynamics of metabolic activity. By fitting a polynomial function to the time-activity curves (TACs), it enables the characterization and subsequent removal of the overall trend in radiotracer uptake over time. The residuals derived from this polynomial fit represent a possibility to obtain inherent physiological fluctuations, facilitating subsequent calculations of inter-regional molecular connectivity.

#### (A2) Spatiotemporal Filter

The spatiotemporal gradient filter, as employed by ^9^, targets short-term fluctuations in glucose uptake. This filter isolates short-term resting-state fluctuations by removing the effect of radiotracer accumulation and low-frequency components of the signal. It effectively adjusts for the mean signal without resorting to global signal regression, thereby avoiding the creation of spurious anticorrelations in the data. The filter utilized a spatial Gaussian standard deviation of one voxel and a temporal Gaussian standard deviation of 2 frames as suggested by Jamadar et al ^9^.

#### (A3) Band-pass Filter

In fMRI connectivity analysis, a band-pass filter is commonly applied to isolate the frequency range corresponding to the resting-state fluctuations of interest. By selectively passing frequencies within this range, the band-pass filter enhances the detection of (presumably) coherent neural activity while suppressing noise and artifacts. Like fMRI resting-state analyses, a band-pass filter can also be applied to fPET TACs. We set the passband frequency for the fPET data to match that used in the fMRI analysis [0.01, 0.1].Hz.

#### (A4) Baseline normalization

This involves a systematic normalization and correlation-based approach. Initially, the entire brain’s mean signal intensity is computed at each time point, serving as a reference for subsequent normalization. Each ROI is then adjusted relative to this global mean value (i.e. dividing a regional TAC by the global TAC at each time point), facilitating comparability across brain regions ^47,48^.

#### (A5) Euclidean distance

In contrast to the approaches above, which necessitates prior filtering to mitigate the influence of baseline uptake, utilizing Euclidean similarity on PET data does not require such preprocessing. Euclidean similarity is based on the Euclidean distance between each pair of time-activity curves (TACs). It is defined as one minus the normalized distance i.e. divided by the maximum distance among pairs of TACs, resulting in values scaled to the range [0, 1]. Due to the heavy-tailed (left-skewed) distribution of Euclidean similarity values, a Fisher z-transformation is applied to normalize the data. Following the transformation, the values are rescaled to the range [0, 1] to maintain consistency with the original scale ^13^.

#### (B1) Covariance matrix

refers to the statistical covariation of radiotracer activity levels across different brain regions over a group of individuals and previously served as a surrogate measure for PET connectivity ^2,3,49^. This approach mostly involves constructing a single image for each participant from the PET data, typically the standard uptake value (SUV). To facilitate inter-subject comparison and mitigate individual variability, these images are typically normalized by the average grey matter value (SUVR), ensuring that differences in global metabolic activity levels are accounted for ^15^.

#### (B2) SICE

Sparse inverse covariance estimation, also known as Gaussian graphical models or graphical Lasso, is a refined method used for estimating molecular covariance. SICE identifies the conditional dependencies between variables in a dataset while promoting sparsity, meaning many entries in the inverse covariance matrix are forced to be zero. This is achieved through regularization techniques that penalize the absolute values of the matrix entries, resulting in a sparse representation that highlights the most significant relationships among variables. This approach is particularly useful in scenarios where the number of subjects included in the analysis is smaller than the number of ROIs, which is valuable in connectivity studies aimed at assessing the whole-brain connectome ^16,26^. In this scenario, we used the graphical Lasso approach with the regularization parameter set to 0.1 and the maximum number of iterations was set to one thousand.

### Statistical comparison

To estimate and visualize the different molecular connectivity or covariance matrices for each method across various levels of network granularity, three functional atlases were selected: the Yeo (17 Network) atlas ^50^ combined with other regions from the Harvard-Oxford atlas distributed with FSL (ROI 18: Striatum & Thalamus; ROI 19: Amygdala & Hippocampus) and the Schaefer 100 and 300 parcellation atlas ^51^, whose networks have been related to the Yeo atlas. Molecular (PET) and functional (fMRI) connectivity was assessed by computing Pearson’s partial correlation of the residual time courses between pairs of brain regions ^52,53^. Head motion was accounted for by incorporating the six realignment parameters as nuisance variables in the partial correlation calculation. We calculated partial correlations for connectivity estimated using the 3^rd^ order polynomial, baseline normalization, spatiotemporal filter for fPET, and band-pass filtering for both fPET and fMRI. For the estimate of molecular covariance metrics, partial correlations were similarly employed, but instead of head motion correction, correlations were adjusted for signal contributions from other brain regions. This approach facilitated the isolation of pairwise information while reducing confounding effects, resulting in a more precise characterization of covariance patterns ^26^.

To aggregate findings across participants, correlation coefficients of all connectivity metrics were averaged after Fisher Z-transformation. Finally, the average Fisher Z-values were inverted back to the correlation values. Next, each method’s average connectivity or covariance matrix was subjected to hierarchical clustering, with chebychev/max distance serving as the distance metric. This approach enabled both visualization and quantification of the distinct differences in patterns generated by each approach. Eigenvalue decomposition was conducted to better assess the spatial structure of each filter’s correlation/covariance matrix, and the results were visualized using a scree plot.

## Experimental Results

### Correlation and Covariance Matrices

The correlation matrices in Figure 3 illustrate distinct molecular connectivity/covariance patterns between the various techniques.

**Figure 3:**
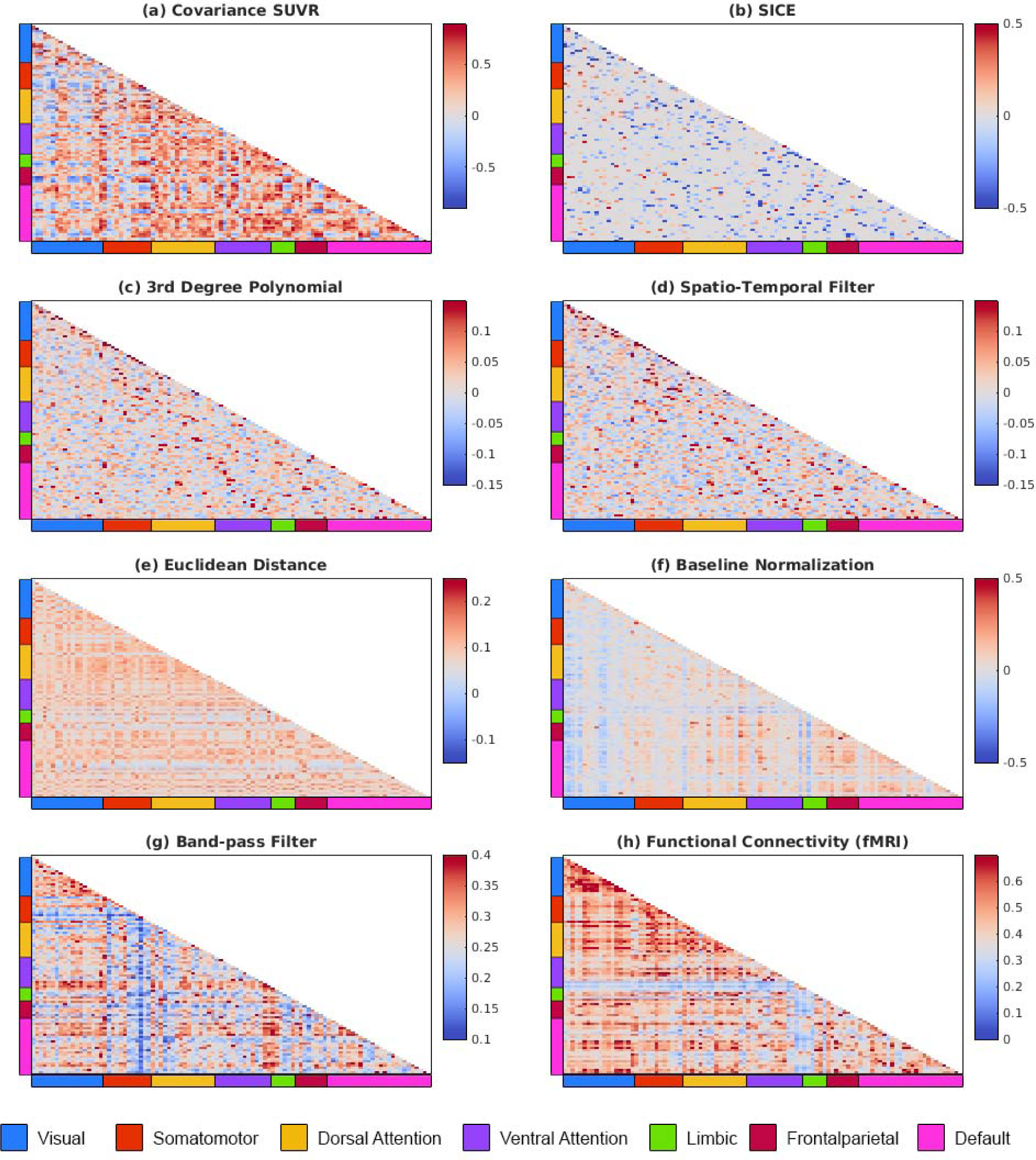
Overview of averaged connectivity/covariance matrices estimated using different techniques divided into regions and fMRI-based networks by the Schaefer 100 atlas. The highest network structure was observed for the band-pass filter, baseline normalization and Euclidian distance metric, whereas the 3^rd^ order polynomial and spatio-temporal filter lacked organizational structure and low correlation values. For the across-subject metrics, the covariance matrix showed higher structure than SICE.

Matrices obtained with the band-pass filter and baseline normalization were characterized by structured network organization and high correlation values indicative of strong inter-regional interactions (Figure 3f-g). However, correlations were markedly influenced by residual baseline radiotracer uptake for the latter approach. This suggests that the normalization does not sufficiently remove this effect (Figure 4d), leading to correlation values primarily driven by baseline uptake rather than moment-to-moment fluctuations. The Euclidian distance metric showed intermediate network structure, most likely due to the similarity of TACs across brain regions (Figure 4a). Compared to fMRI-based functional connectivity, the band-pass filter showed network structure to a similar extent but different organizational pattern, highlighting the difference between the various imaging modalities (Figure 3 and 6g-h). In contrast, the 3^rd^-order polynomial and spatial-temporal filter methods yield matrices almost devoid of network structure (Figure 3 and 6c-d), attributed to low correlation values, which in turn emerged from high levels of noise in the signal (Figure 4b-c). These methods operate like high-pass filters, effectively eliminating low-frequency baseline tracer uptake (Figure 4). However, the band-pass filter successfully removes high-frequency noise, yielding the highest molecular connectivity values. Among the across-subject molecular covariance and SICE methods, both exhibited high correlation values, but the latter showed less network organization, (Figure 3 and 6a-b).

**Figure 4:**
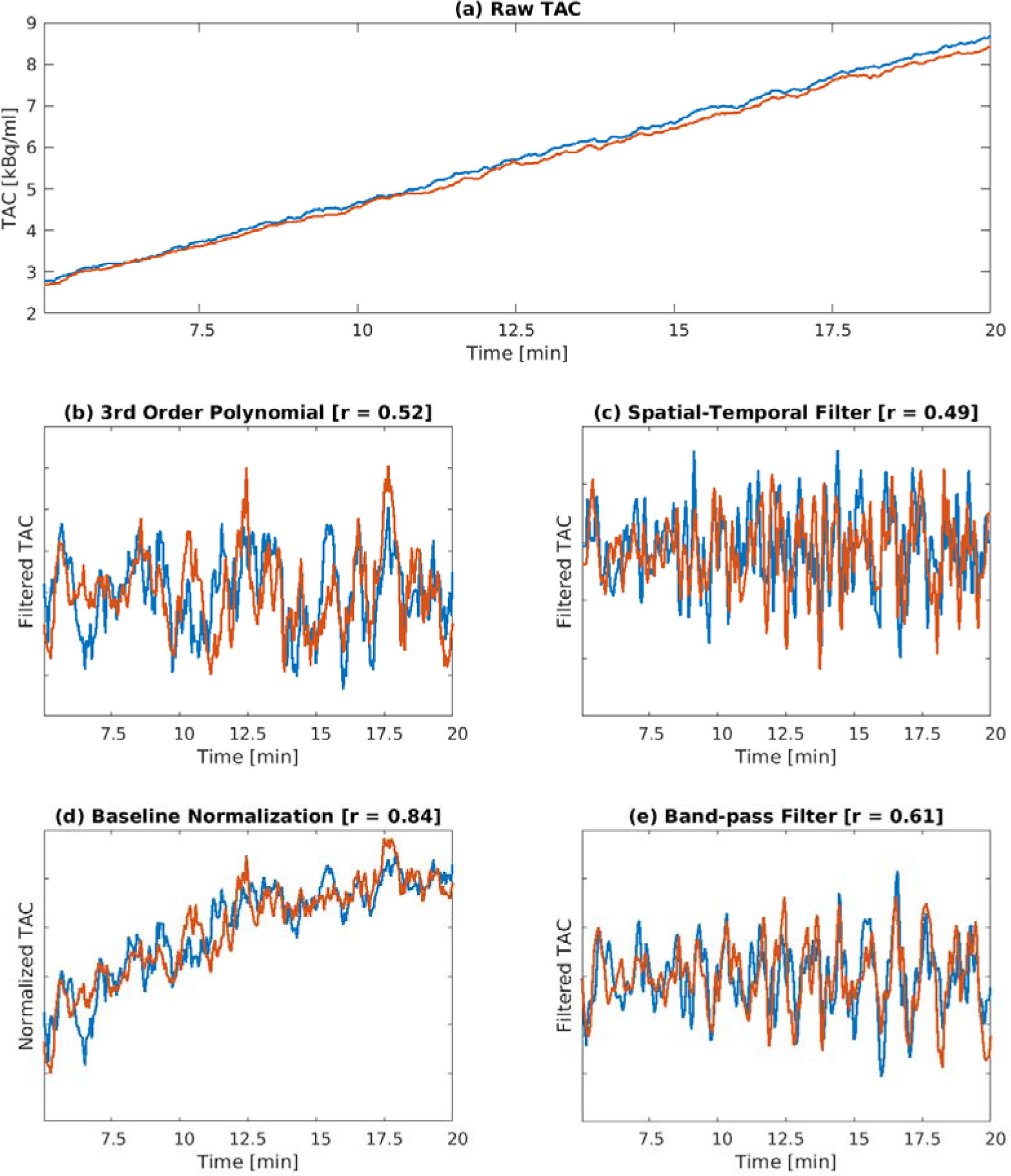
Time course of two brain regions from the frontoparietal network from a representative subject before and after the application of each filtering technique. The raw TACs (a) are used as inputs for the Euclidian distance metric. All other approaches aim to compute connectivity by correlation of moment-to-moment fluctuations in the signal. While the 3^rd^ order polynomial (b) and spatio-temporal filter (c) were able to remove the baseline radiotracer uptake, this was not the case for the baseline normalization approach (d), demonstrating that residual baseline radiotracer uptake (instead of moment-to-moment fluctuations) drives the correlations. Still, signals in b and c were characterized by high noise levels, which were effectively removed by the band-pass filter (e).

**Figure 5:**
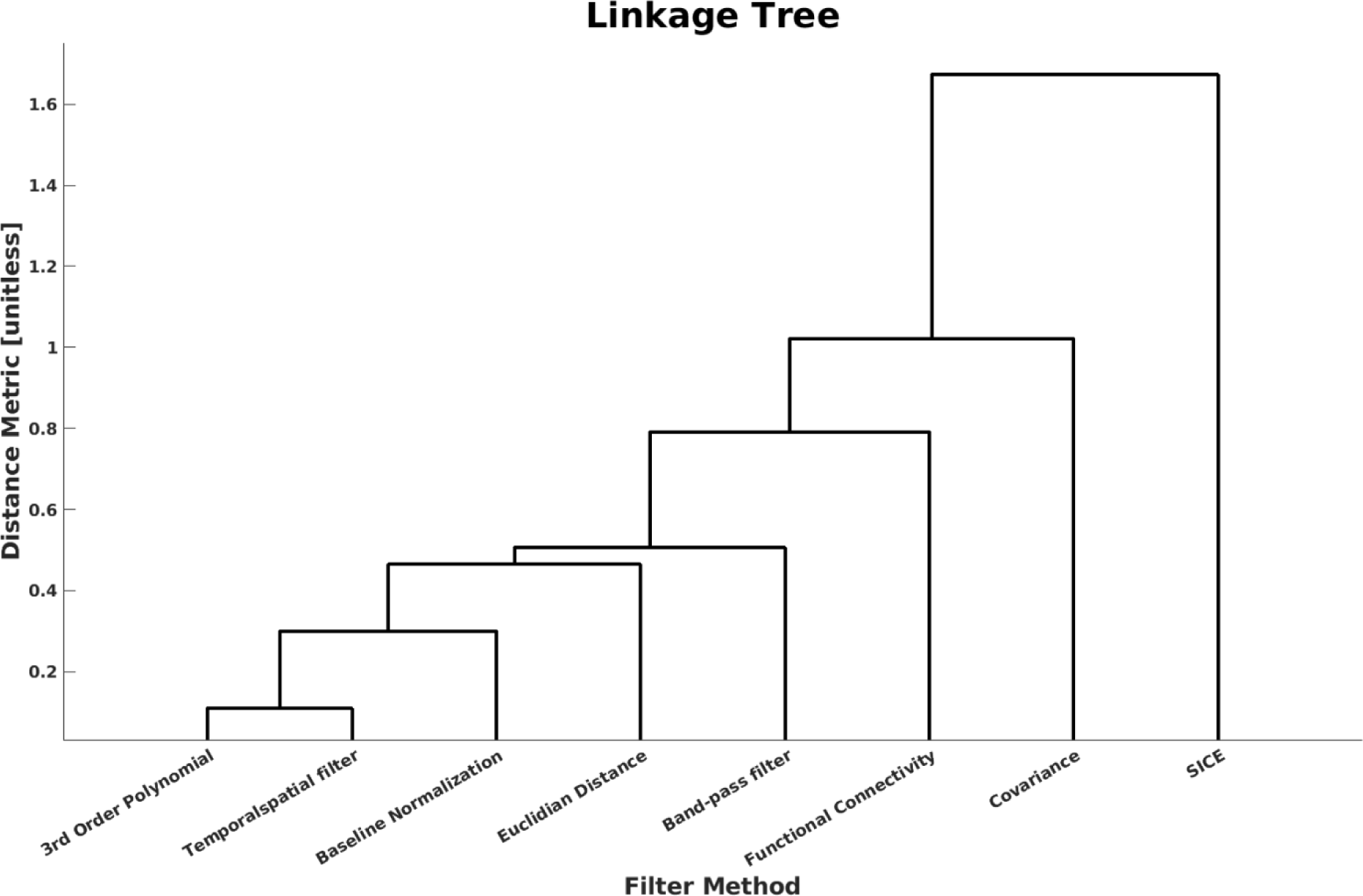
Data-driven clustering of the different connectivity/covariance metrics from the Schaefer 300 Network atlas. Most of the within-subject techniques grouped together, in particular the 3^rd^ order polynomial, spatio-temporal filter and band-pass filter, which is evident since the former two act as high-pass filters.

**Figure 6:**
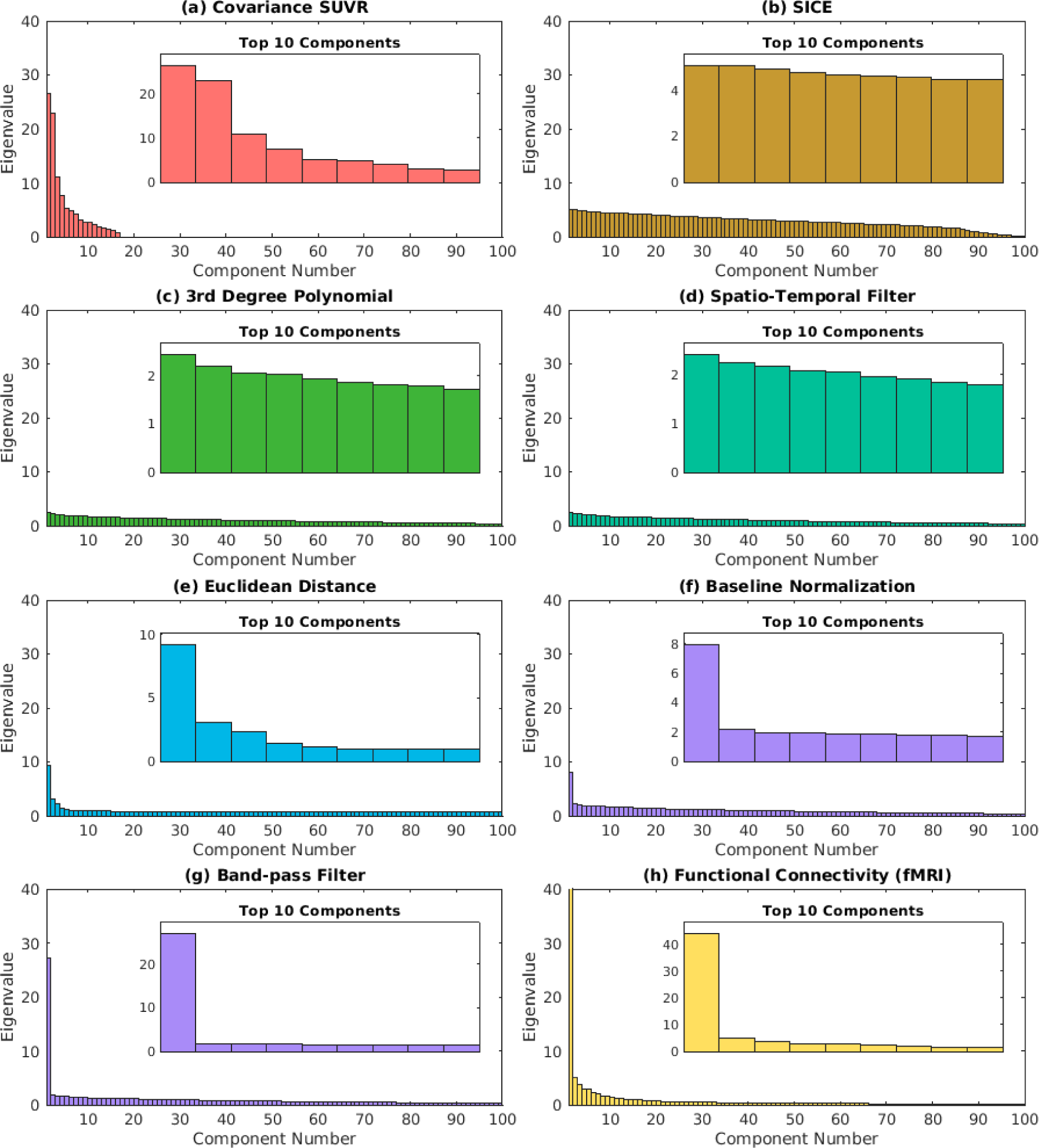
Eigenvalue decomposition of the averaged connectivty/covariance matrices estimated using each technique divided into regions and fMRI-based networks by the Schaefer 100 atlas. A high eigenvalue component suggests the presence of strong underlying structures or patterns within the data, which can be seen in the covariance SUVR, band-pass and functional connectivity methods and, to a lesser extent, euclidean distance and baseline normalization. Conversely, the absence of clear peaks suggests a correlation matrix with weak or random associations among variables, implying a less organized or structured dataset, as seen in SICE, 3rd degree polynomial and spatio-tempral filter.

### Hierarchical Clustering and eigenvalue decomposition

Hierarchical clustering provides a data-driven approach to discern differences among techniques. As depicted in Figure 4, the linkage tree highlights the dissimilarity of across-subject methods such as SICE and molecular covariance from others. The 3^rd^ order polynomial, temporal-spatial, and band-pass filter methods cluster closely together, with baseline normalization and functional connectivity grouped in proximity. This observation did not change considerably with the number of brain regions (Supplementary Figure 1).

Eigenvalue decomposition correlations’ and covariance matrices’ network organization (Figure 3). The results confirmed that the band-pass filter and fMRI functional connectivity exhibited significantly higher loading on the first eigenvalue than others (Figure 6g-h). Across-subject molecular covariance, Euclidean distance, and baseline normalization displayed a reduced organization (Figure 6a, e-f). In contrast, SICE, 3rd degree polynomial, and spatio-temporal filters showed low eigenvalue loadings, suggesting low network organization (Figure 6b-d).

These findings underscore the nuanced differences across various approaches at high-temporal resolution, highlighting the importance of methodological considerations when interpreting molecular connectivity estimates.

## Discussion of Experimental data

The experimental work supports the standardized nomenclature for commonly employed molecular connectivity and covariance techniques based on available literature, aiming to provide a clearer understanding of their practical application. Using high-temporal resolution [^18^F]FDG fPET data, this data-driven approach represents a showcase of the feasibility for deriving molecular connectivity and covariance using common techniques. Thus, the combination of literature-based nomenclature with a data-driven approach provides a robust and comprehensive framework.

Computing associations of moment-to-moment fluctuations in the metabolic signal using a band-pass filter showed a similar level of network organization but a different pattern to rsfMRI functional connectivity. In contrast, other methodological approaches require further development to obtain robust molecular connectivity through more accurately elimination of baseline uptake (baseline normalization approach) or high frequency noise (3^rd^ order polynomial and spatio-temporal filter, Figure 3 and 4). However, studies have demonstrated the efficacy of the latter two at lower temporal resolutions, such as 16 seconds ^9^ or 1 minute ^15^. At a high temporal resolution, these issues can be overcome e.g., by a band-pass filter, which removes both low-frequency signal changes in radiotracer uptake (such as irreversible binding for [^18^F]FDG) as well as high-frequency noise (Figure 4). Furthermore, the 3^rd^ order polynomial filter approach was able to eliminate the baseline uptake, making the computation of task effects feasible ^4,41,54^. Of note, when using irreversibly binding radioligands, such as [^18^F]FDG, a bolus + infusion protocol allows for a more accurate computation of molecular connectivity from moment-to-moment signal fluctuations, since free radioligand is constantly provided throughout the experiment.

On the other hand, associations in radiotracer kinetics are unique to PET imaging, showing a distinct kinetic connectivity structure across the brain. Conversely, the computation of covariance matrices unveils reduced network organization. Particularly, SICE yields a sparse covariance matrix, prioritizing the identification of conditional dependencies among variables. This emphasis on uncovering direct relationships between variables proves invaluable, especially in scenarios where understanding the relations within a specific cohort is paramount.

Without dynamic PET data, molecular covariance can be estimated. This method holds promise for its straightforward application and data acquisition within clinical settings, (only requiring a static image per subject). However, covariance metrics are not without their limitations, e.g., when conducting a connectome-wide investigation, where the number of ROIs surpasses the number of subjects and relatively small sample sizes may introduce a potential bias, which are commonplace in PET studies. Recently, it has been suggested that partial correlation analysis should be used, overcoming the constraints of simple correlation analysis, which solely captures pairwise information and fails to characterize the effects of multiple brain regions interacting collectively ^26^. A more advanced approach, SICE, has been advocated to address this concern. Our findings reveal that while molecular covariance and SICE methods use the same data, SICE diverges from molecular connectivity metrics and its covariance counterpart. While covariance metrics primarily assess the relationships between variables, reflecting their co-variation, SICE delves into conditional dependencies. This emphasis renders SICE particularly adept at uncovering nuanced associations within complex datasets, focusing on only the strongest association within a group.

In juxtaposing molecular connectivity with its fMRI counterpart functional connectivity, it becomes evident that both methods yield divergent connectivity measures and can offer complementary perspectives on brain connectivity, accentuating the importance of multimodal approaches to connectivity studies. Thus, the integration of individual-level fMRI and fPET connectivity presents intriguing avenues for investigating brain function across various organizational levels, given the different underlying basis of BOLD fMRI functional connectivity (blood flow and oxygenation) and [^18^F]FDG fPET molecular connectivity (glucose metabolism), which are connected through overlapping neurophysiological effects (glutamate and GABA signaling) ^4–7,41^.

### Limitations, Outlook, and Conclusion

This work does not aim to provide an in-depth comparison of all available techniques or determination of optimal parameter settings for each method. This exceeds the scope of the current manuscript but on the other hand would hardly affect the terminology. The main goal was to define a common nomenclature for the available techniques, drawing from both literature and empirical data perspectives. More specifically, this work highlights the different opportunities to derive individual-level molecular connectivity and kinetic connectivity as well as group-level molecular covariance from PET data. We would like to emphasize that all these approaches are valid and valuable but require distinct terminology due to the underlying differences in the assumptions, calculations, and outcome metrics. We also acknowledge that PET-based connectivity/covariance and fMRI functional connectivity were obtained from different subjects. As the evaluation of moment-to-moment fluctuations in PET signals is still in its infancy, future work should aim to identify the underlying neurophysiological mechanisms. This will boost the interpretation of PET-based connectivity approaches and may aid in the identification of pathophysiological processes in brain disorders.

Investigating the differences in brain network interactions based on different imaging techniques represents a promising opportunity for future work. While, exploring alternative tracers and more advanced methodologies, such as directional molecular connectivity, holds great promise for advancing knowledge of brain networks, this progress requires a clear terminology for the distinct types of brain connectivity.

## Acknowledgements

We would like to thank the study staff of the Neuroimaging Lab at the Department of Psychiatry and Psychotherapy for clinical, scientific, technical and administrative support. We are especially grateful to .M. Klöbl, L. Rischka, GM James, T. Vanicek, J. Unterholzner, P. Michenthaler, A. Kautzky, M. Hienert, T. Balber, V. Pichler, E. Petronas, M. Hartenbach, E. Winkler-Pjrek, W. Wadsak, M. Mitterhauser and S. Kasper for all their support.

## Data Availability Statement

Raw data will not be made publicly available due to reasons of data protection. Processed data and custom code can be obtained from the corresponding author with a data-sharing agreement, approved by the departments of legal affairs and data clearing of the Medical University of Vienna.

## Supplementary Figure and Table

**Supplementary figure 1:**
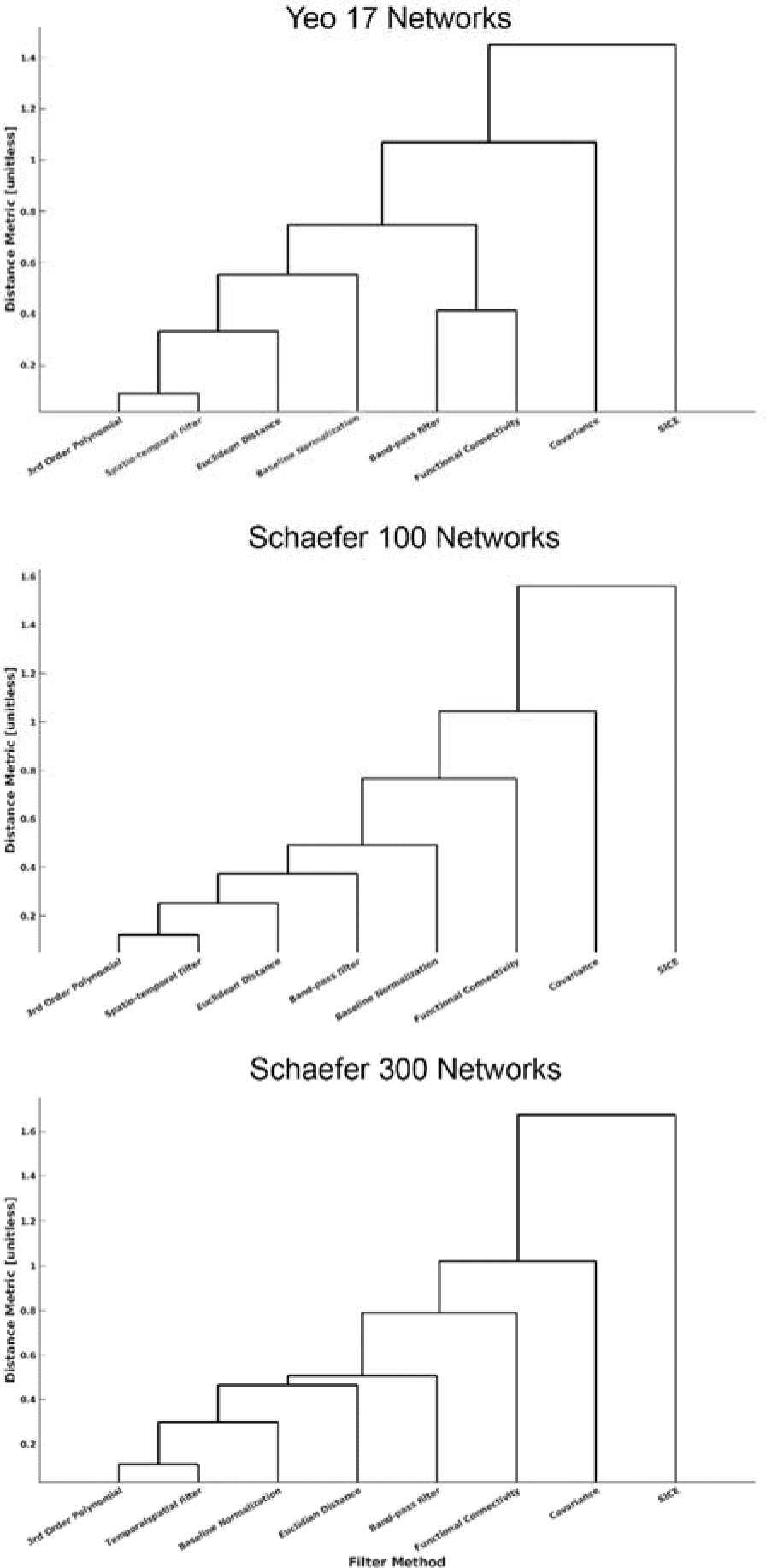
Data-driven clustering performed on different connectivity/covariance metrics derived from the (a) Yeo 17 Networks, (b) Schaefer 100, and (c) Schaefer 300 Network atlases. Notably, regardless of network granularity (i.e., the number of ROIs), the clustered techniques demonstrated stability. The majority of within-subject techniques, notably the 3^rd^ order polynomial, spatio-temporal filter, and band-pass filter, remained closely grouped together. In contrast, covariance and SICE metrics exhibited distinct clustering, indicating greater dissimilarity from other methods in terms of connectivity patterns.

**Supplementary Table 1:** Exemplary overview of proposed terms for PET connectivity and covariance. terms based on radioligands. A naming convention is proposed for additional radiotracers, wherein the convention involves stating the target (prefix) followed by the utilized connectivity or covariance. It is important to note that the feasibility of estimating molecular connectivity (i.e., moment-to-moment fluctuations) for all radiotracers in this list is not yet established (marked italics).

| Tracer | Molecular connectivity (MC) term | Kinetic connectivity (KC) term | Molecular covariance (MCov) term |
| --- | --- | --- | --- |
| [<sup>18</sup>F]FDG | Metabolic connectivity (M-MC) | Metabolic kinetic connectivity (M-KC) | Metabolic covariance (M-MCov) |
| [<sup>15</sup>O]H<sub>2</sub>O | Cerebral blood flow connectivity (CBF-MC) | Cerebral blood flow kinetic connectivity (CBF-KC) | Cerebral blood flow covariance (CBF-MCov) |
| [<sup>18</sup>F]FDOPA | Dopamine synthesis connectivity (DAS-MC) | Dopamine synthesis kinetic connectivity (DAS-KC) | Dopamine synthesis covariance (DAS-MCov) |
| [*carbonyl*-<sup>11</sup>C]WAY100635 | *5-HT<sub>1A</sub> connectivity (5-HT<sub>1A</sub>-MC)* | 5-HT<sub>1A</sub> kinetic connectivity (5-HT<sub>1A</sub>-KC) | 5-HT<sub>1A</sub> covariance (5-HT<sub>1A</sub>-MCov) |
| [<sup>11</sup>C]DASB | *SERT connectivity (SERT-MC)* | SERT kinetic connectivity (SERT-KC) | SERT covariance (SERT-MCov) |
| [<sup>18</sup>F]flutemetamol | *Amyloid beta connectivity (Abeta-MC)* | Amyloid beta kinetic connectivity (Abeta-KC) | Amyloid beta covariance (Abeta-MCov) |
| [<sup>18</sup>F]flortaucipir | *Tau connectivity (tau-MC)* | Tau kinetic connectivity (tau-KC) | Tau covariance (tau-MCov) |

